# Reclassification of *Shigella* species as later heterotypic synonyms of *Escherichia coli* in the Genome Taxonomy Database

**DOI:** 10.1101/2021.09.22.461432

**Authors:** Donovan H. Parks, Maria Chuvochina, Peter R. Reeves, Scott A. Beatson, Philip Hugenholtz

## Abstract

Members of the genus *Shigella* have high genomic similarity to *Escherichia coli* and are often considered to be atypical members of this species. In an attempt to retain *Shigella* species as recognizable entities, they were reclassified as *Escherichia* species in the Genome Taxonomy Database (GTDB) using an operational average nucleotide identity (ANI)-based approach nucleated around type strains. This resulted in nearly 80% of *E. coli* genomes being reclassified to new species including the common laboratory strain *E. coli* K-12 (to ‘*E. flexneri’*) because it is more closely related to the type strain of *Shigella flexneri* than it is to the type strain of *E. coli*. Here we resolve this conundrum by treating *Shigella* species as later heterotypic synonyms of *E. coli*, present evidence supporting this reclassification, and show that assigning *E. coli*/*Shigella* strains to a single species is congruent with the GTDB-adopted genomic species definition.

## Introduction

The genus *Escherichia* currently comprises the six species *E. coli* (Castellani and Chalmers, 1919), *E. hermannii* (Brenner et al., 1982), *E. fergusonii* (Farmer et al., 1985), *E. albertii* (Huys et al., 2003), *E. marmotae* (Liu et al., 2015), and the recently described *E. ruysiae* (van der Putten et al., 2021). However, a recent phylogenetic study places *E. hermannii* within the effectively published but currently unvalidated genus ‘*Atlantibacter’* (Hata et al., 2016), and both the NCBI (Schoch et al., 2020) and GTDB (Parks et al., 2018) taxonomies follow this reclassification. There are also a number of strains that have been recognized as distinct *Escherichia* lineages that have no assigned species name and are referred to as ‘cryptic clades’ (Walk et al., 2009). However, the most enduring anomaly with this genus is paraphyly between *E. coli* and the four currently recognized *Shigella* species (The et al. 2016; Pettengill et al., 2016; Hu et al., 2019), which has been attributed to independent acquisition of the pINV virulence plasmid and subsequent niche adaptation (Pupo et al., 2000; Lan and Reeves 2002; Yang et al., 2005).

Numerous studies have shown that *E. coli* and *Shigella* species have high genomic similarity (Brenner et al., 1973; Richter and Rosselló-Móra 2009; Jain et al., 2018; Ciufo et al., 2018). Consequently, it is widely recognized that *E. coli* and *Shigella* species could be considered a single species (Lan and Reeves, 2002; Richter and Rosselló-Móra, 2009; van den Beld and Reubsaet, 2012); however, this reclassification has not been adopted due to the medical significance of *Shigella* as pathogens causing shigellosis, a form of bacillary dysentery (Ewing 1949; Sahl et al., 2015). At the same time, enteroinvasive *E. coli* (EIEC) also causes dysentery making the clinical distinction between *E. coli* and *Shigella* ambiguous as few biochemical properties distinguish *Shigella* from EIEC (Johnson 2000; Yang et al., 2005; van den Beld 2012; Hendriks et al., 2020). Studies have also shown that *E. coli* and *Shigella* species are paraphyletic and that *S. boydii, S. dysenteriae*, and *S. flexneri*, primarily defined based on serotype, are not monophyletic entities (Sahl et al., 2015; Pettengill et al., 2016; Pupo et al., 2000; Hu et al., 2019) further justifying the reclassification of *Shigella* species as *E. coli*.

The Genome Taxonomy Database (GTDB) has adopted a genomic species definition based on average nucleotide identity (ANI) and alignment fraction (AF) nucleated around genomes from type strains for delineating species (Parks et al., 2020) and an estimation of time of divergence for circumscribing higher taxonomic ranks (Parks et al., 2018). Application of these criteria resulted in *Shigella* species being reassigned to the genus *Escherichia* as ‘*E. flexneri’* and ‘*E. dysenteriae’* with *S. sonnei* and *S. boydii* being classified as synonyms of ‘*E. flexneri’* due to their high genomic relatedness (>97%). Notably, nearly 80% of the genomes in GTDB R06-RS202 formerly classified as *E. coli* were reclassified as either ‘*E. flexneri’* or ‘*E. dysenteriae’* as a result of the adopted criteria for delineating species. This includes the common laboratory strain *E. coli* str. K-12 which was assigned to ‘*E. flexneri’* due to its higher genomic similarity to the type strain of ‘*E. flexneri’* than to the type strain of *E. coli* (U5/41 = ATCC 11775; Meier-Kolthoff et al., 2014).

A practical consequence of these GTDB reassignments is that traditional properties of species such as ‘*E. dysenteriae’* and ‘*E. flexneri’* being comprised of human pathogenic strains are no longer applicable. As this is likely to result in confusion, we propose that *Shigella spp*. be treated as later heterotypic synonyms of *E. coli* in GTDB. Here we put forth the evidence supporting this reclassification in anticipation of introducing this change in the next release of GTDB.

## Results and discussion

### Reclassification of *Escherichia coli* and *Shigella* in GTDB

GTDB assigns strains to species based on the ANI and AF to genomes assembled from the type strain of the species (henceforth referred to as type strain genomes) or to a selected representative genome of the species when a type strain genome is not available (Parks et al., 2020). This provides quantitative criteria for establishing species membership that is consistent with the majority of species assignments defined using a polyphasic approach. In GTDB release R06-RS202, 68.2% of genomes with an NCBI species assignment have the same species assignment in the GTDB. Genomes with incongruent GTDB and NCBI species assignments result from the transfer of species to other genera (10.4%), defining overclassified species as synonyms (1.4%), dividing species considered to be underclassified into multiple species (5.9%), and otherwise reclassifying genomes consider misclassified according to the GTDB (14.1%; **Supp. Table 1**). Notably, >50% of incongruent species assignments arise from just nine NCBI species with *E. coli* and *Shigella spp*. being the most conspicuous accounting for 32.6% of all incongruent assignments between GTDB and NCBI (**Table 1**; **Supp. Table 2**). This is in contrast to other *Escherichia* species where all or nearly all genomes assigned to these species within GTDB have the same assignment in the NCBI Taxonomy (**Table 1**).

**Table 1.** GTDB R06-RS202 assignment of genomes classified as *Escherichia* or *Shigella* within the NCBI taxonomy.

| NCBI species | No. genomes | No. reassigned genomes | Reassigned genomes (%) | GTDB species |
| --- | --- | --- | --- | --- |
| *Escherichia coli* | 20,973 | 16,609 (79.19%) | 28.28 | *E. flexneri*: 12,400 (59.12%); *E. coli*: 4,364 (20.81%); *E. coli*_D: 2,094 (9.98%); *E. dysenteriae*: 2,044 (9.75%); *E. coli*_C: 45 (0.21%); *E. sp000208585*: 9 (0.04%); *E. sp005843885*: 2 (0.01%); *Klebsiella quasipneumoniae*: 1 (0.00%); *E. coli*_E: 1 (0.00%); *Phytobacter ursingii*: 1 (0.00%); *Citrobacter freundii*: 1 (0.00%); *Enterobacter roggenkampii*: 1 (0.00%); *Pseudomonas*_E *sp002113295*: 1 (0.00%); *E. albertii*: 1 (0.00%); *Kluyvera georgiana*_A: 1 (0.00%); *Hafnia alvei*: 1 (0.00%); *E. marmotae*: 1 (0.00%); *Citrobacter gillenii*: 1 (0.00%); *Providencia rettgeri*_D: 1 (0.00%); *Enterobacter hormaechei*_A: 1 (0.00%); *Klebsiella aerogenes*: 1 (0.00%); *Klebsiella variicola*: 1 (0.00%) |
| *Shigella sonnei* | 1,368 | 1,368 (100.00%) | 2.33 | *E. flexneri*: 1,354 (98.98%); *E. coli*_D: 13 (0.95%); *Serratia marcescens*_K: 1 (0.07%) |
| *Shigella flexneri* | 706 | 706 (100.00%) | 1.20 | *E. flexneri*: 705 (99.86%); *E. coli*: 1 (0.14%) |
| *Shigella boydii* | 120 | 120 (100.00%) | 0.20 | *E. flexneri*: 111 (92.50%); *E. coli*_D: 5 (4.17%); *Serratia marcescens*_I: 3 (2.50%); *E. dysenteriae*: 1 (0.83%) |
| *Shigella dysenteriae* | 60 | 60 (100.00%) | 0.10 | *E. flexneri*: 41 (68.33%); *E. dysenteriae*: 18 (30.00%); *E. coli*_D: 1 (1.67%) |
| *Escherichia fergusonii* | 74 | 1 (1.35%) | 0.00 | *E. fergusonii*: 73 (98.65%); *E. flexneri*: 1 (1.35%) |
| *Escherichia albertii* | 98 | 0 (0.00%) | 0.00 | *E. albertii*: 98 (100.00%) |
| *Escherichia marmotae* | 34 | 0 (0.00%) | 0.00 | *E. marmotae*: 34 (100.00%) |
| *Escherichia ruysiae* | n/a | n/a | n/a | (proposed after GTDB R06-RS202; see *Methods*) |
| Unclassified *Escherichia* sp. | 153 | 153 (100.00%) | 0.26 | *E. marmotae*: 42 (27.45%); *E. sp005843885*: 35 (22.88%); *E. sp000208585*: 22 (14.38%); *E. flexneri*: 18 (11.76%); *E. coli*_C: 17 (11.11%); *E. coli*_D: 8 (5.23%); *E. coli*: 3 (1.96%); *E. sp004211955*: 2 (1.31%); *E. sp001660175*: 2 (1.31%); *Citrobacter freundii*: 1 (0.65%); *E. sp002965065*: 1 (0.65%); *PseudE. vulneris*: 1 (0.65%); *PseudE. sp002918705*: 1 (0.65%) |
| Unclassified *Shigella* sp. | 112 | 112 (100.00%) | 0.19 | *E. flexneri*: 107 (95.54%); *Proteus* *sp001722135*: 2 (1.79%); *E. coli*_D: 2 (1.79%); *E. coli*: 1 (0.89%) |

Strains have traditionally been assigned to *E. coli* and *Shigella spp*. based on biochemistry and serotyping (Pettengill et al., 2016; Chattaway et al., 2017) which is in contrast to the genomic species definition (Doolittle and Papke 2006; Konstantinidis et al., 2006; Richter and Rosselló-Móra, 2009) adopted by GTDB. Specifically, GTDB delineates species using a fixed AF threshold of 0.65 and an ANI threshold that is allowed to vary between 95% and 97% in order to preserve the majority of existing species classifications (Parks et al., 2020). Despite using a flexible ANI threshold, ‘*E. sonnei’* and ‘*E. boydii’* are considered later heterotypic synonyms of ‘*E. flexneri’* in GTDB as the ANI between these type strain genomes is 97.9% and 97.8%, respectively (**Fig. 1**). Furthermore, 79.2% of the 20,973 genomes classified as *E. coli* at NCBI are reassigned within GTDB with the majority being reclassified as ‘*E. flexneri’* (12,400 genomes) or ‘*E. dysenteriae’* (2,044 genomes; **Table 1**). Similarly, the majority (68.3%) of genomes classified as *S. dysenteriae* at NCBI are classified as ‘*E. flexneri’* in GTDB (**Table 1**). GTDB species assignments reflect the closest type strain genome as determined using ANI and this large number of reassignments illustrate the degree to which the traditional classification of *E. coli* and *Shigella spp*. conflict with the adopted genomic species definition (**Fig. 1**). A few clear misclassifications within the NCBI taxonomy are also evident such as the reassignment of a *S. sonnei* and three *S. boydii* genomes to species in the genus *Serratia* in GTDB (**Supp. Table 3**).

**Figure 1.**
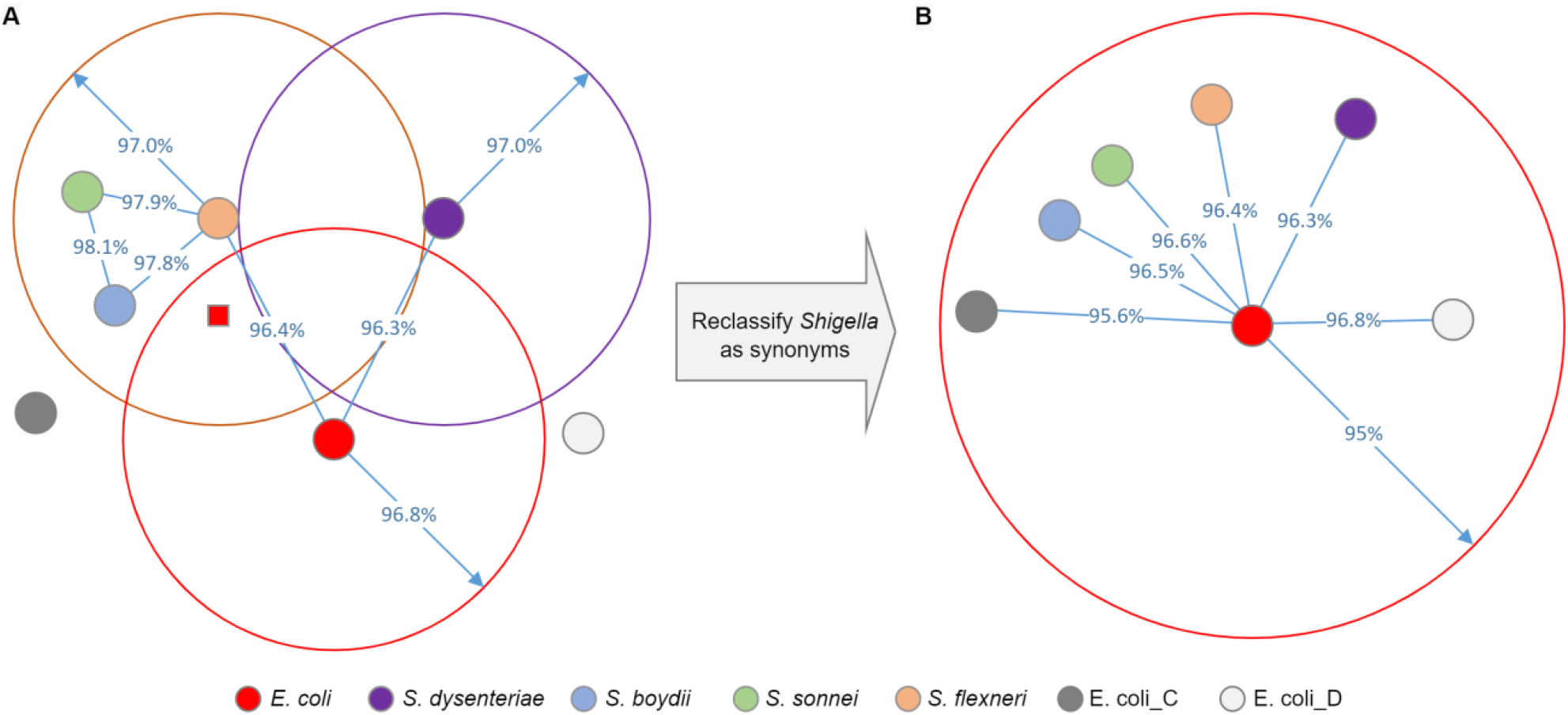
Conceptual diagram showing the high genomic similarity between *Escherichia* and *Shigella* species (**A**), and the proposal to reclassify *Shigella spp*. as later heterotypic synonyms of *E. coli* (**B**). Filled circles indicate genomes assembled from the type strain of *E. coli* or a *Shigella spp*., and the representative genomes from the GTDB species E. coli_C and E. coli_D. The distance between genomes indicates their ANI though the diagram is conceptual as it is not possible to accurately represent the pairwise distance between all genomes. The larger circles depict the ANI criteria used by GTDB for assigning genomes to each species (Parks et al., 2020). This criterion is typically 95% in the GTDB, but the high similarity of these species results in this criterion being increased. The single red square represents a genome traditionally classified as *E. coli* which is reclassified as *S. flexneri* in GTDB since it resides within the ANI threshold of both species, but is more similar to the *S. flexneri* type strain genome. Reclassification of *Shigella spp*. as later heterotypic synonyms of *E. coli* results in a species with a 95% ANI criterion commensurate with the majority of GTDB species. E. coli_C and E. coli_D are also reclassified as *E. coli* as they have an ANI >95% to the type strain of *E. coli*.

### Genomes assembled from type strains of a species are highly similar

Here we verify that type strain genomes from *Escherichia* and *Shigella* species are highly similar to each other giving confidence that the assemblies are of high quality and unlikely to be misclassified (**Table 2**). All species except *E. albertii, E. ruysiae*, and *S. flexneri* had two or more type strain genomes available, and intraspecific type strain genomes had an ANI >99.8% with an AF generally >0.97. Lower AFs were observed in a few cases, but this can be attributed to either assembly quality (e.g., *E. marmotae* assembly GCF_000807695.1 consists of 218 contigs compared to the single chromosome and two plasmids of the *E. marmotae* assembly GCF_002900365.1) or naturally occurring differences in the phage or genomic islands within individual genomes.

**Table 2.**
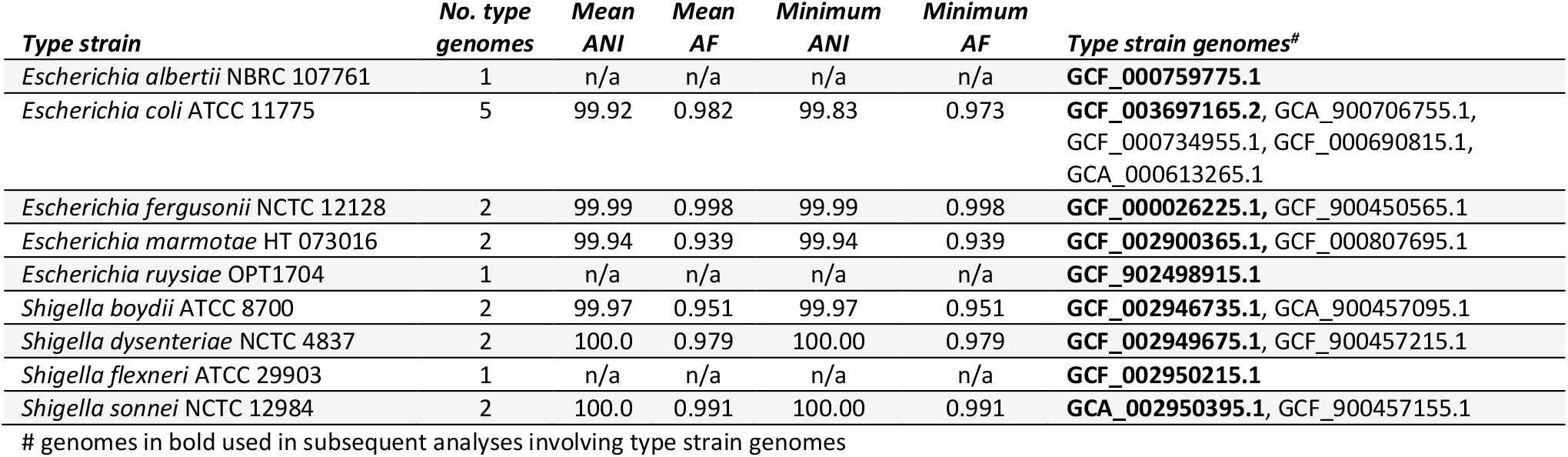
Comparison of genomes assembled from type strains of *Escherichia* and *Shigella* species. # genomes in bold used in subsequent analyses involving type strain genomes

### Phylogeny and genomic similarity of *Escherichia* and *Shigella* type strains

A maximum-likelihood tree was used to establish the phylogenetic relationships between type strain genomes for *Escherichia* and *Shigella* species along with the representative genomes of the seven GTDB R06-RS202 placeholder species within *Escherichia* (**Fig. 2A**; **Supp. Table 4**). As expected, *E. coli* and *Shigella spp*. form a monophyletic lineage which also includes the GTDB species E. coli_D. Genomic species are generally defined as strains with an ANI greater than 94% to 96% and an AF greater than 0.5 to 0.65 (Kostantinidis and Tiedje, 2005; Goris et al., 2007; Varghese et al., 2015; Ciufo et al., 2018; Jain et al., 2019). The type strain genomes for *E. coli* and the *Shigella spp*. along with the representative genomes for E. coli_D and E. coli_C satisfy these criteria with the most dissimilar species being *S. dysenteriae* and E. coli_C at 95.3% ANI and 0.81 AF (**Fig. 2B**; **Supp. Table 5**). Furthermore, all genomes classified as *E. coli*, ‘*E. flexneri’*, ‘*E. dysenteriae’*, E. coli_C, or E. coli_D in GTDB have an ANI ≥95% and AF ≥0.65 to the *E. coli* type strain (ATCC 11775^T^) genome when using FastANI (**Figs. 2C** and **2D**), and thus satisfy the GTDB criteria for classifying these strains as a single species (Parks et al., 2020). This result is robust to the method used to establish genomic similarity as indicated by BLAST-based ANI (Rodriguez-R and Konstantinidis, 2016) providing highly correlated results with all *E. coli*, ‘*E. flexneri’*, ‘*E. dysenteriae’*, E. coli_C, and E. coli_D genomes still reported as having ≥95% ANI to *E. coli* ATCC 11775^T^ (**Supp. Fig. 1**).

**Figure 2.**
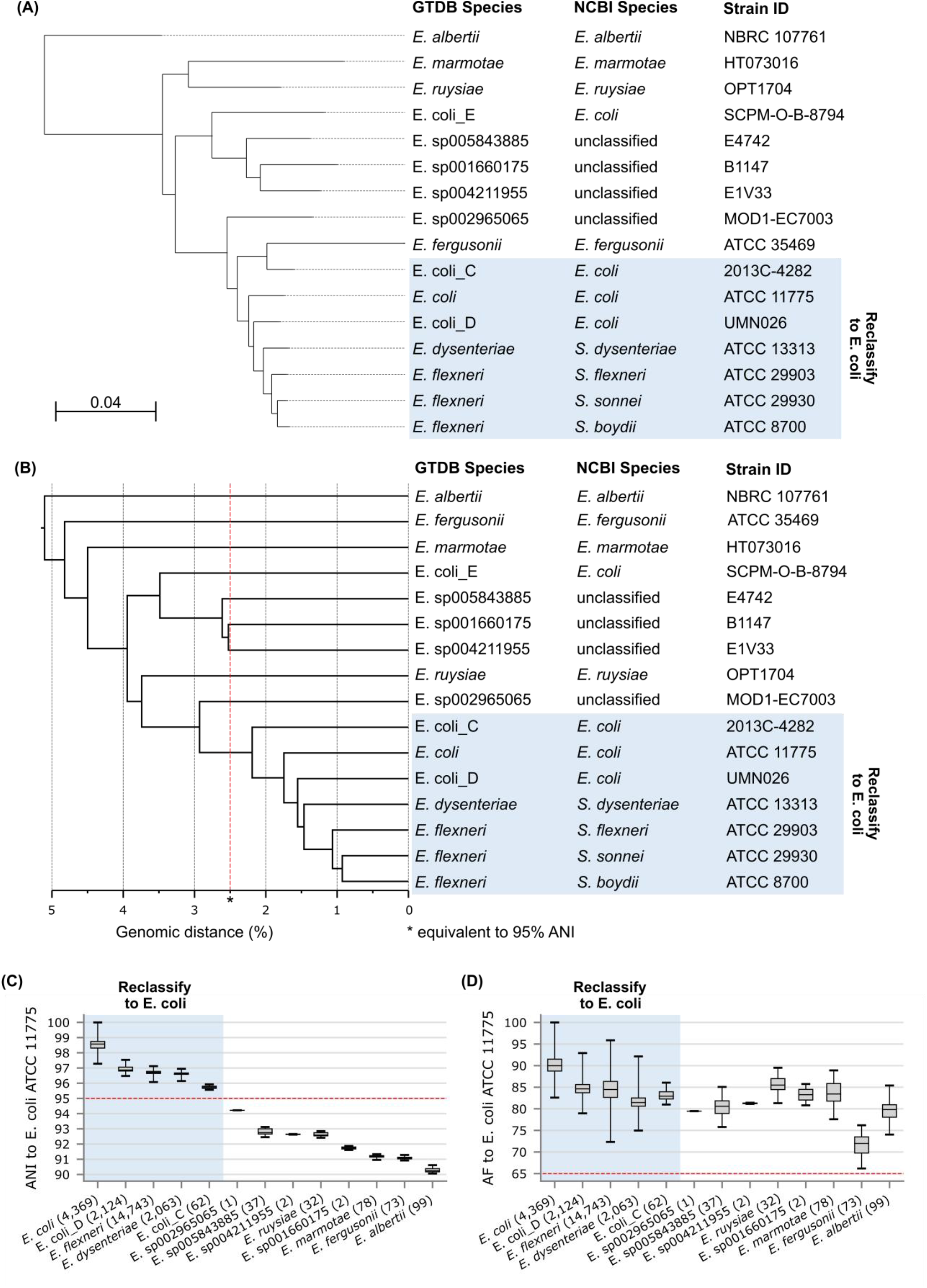
Phylogenetic and genomic similarity of *Escherichia* and *Shigella* species. (**A**) Phylogenetic tree inferred from the concatenated multiple sequence alignment of 2,329 core genes using IQ-Tree under the GTR+F+R5 model. *E. albertii* NBRC 107761^T^ was used to root the tree (*see Methods*). SH-aLRT and UFBoot support values were 100% for all nodes. (**B**) UPGMA tree indicating the genomic distance, i.e. 100-ANI, between *Escherichia* and *Shigella* type strain genomes as determined with FastANI. (**C**) ANI between the *E. coli* ATCC 11775^T^ genome and all genomes classified as *Escherichia* in GTDB R06-RS202. (**D**) AF between the *E. coli* ATCC 11775^T^ genome and all genomes classified as *Escherichia* in GTDB R06-RS202. Box-and-whisker plots show the lower and upper quartiles as a box, the median value as a line within the box, and the minimum and maximum values as whiskers.

### *Escherichia coli* and *Shigella spp*. form a single, distinct ANI species cluster

It is informative to consider the clustering which occurs when considering all pairwise ANI values between *Escherichia* and *Shigella* genomes and not just the ANI to the type strain/representative genomes of species. Here we explore this using Mash which provides a computationally efficient approximation to ANI allowing it to be applied to large numbers of genomes. Specifically, we use Mash to dereplicate all 23,538 high-quality *Escherichia* genomes in GTDB R06-R202 to 1,148 genomes which have an ANI <99% and can be seen as operationally defined strains (*see Methods*). Visualization of these 1,148 operational strains using a 95% ANI clustering criterion indicates that *E. coli*, ‘*E. flexneri’*, ‘*E. dysenteriae’*, E. coli_C, and E. coli_D form a single cluster in support of reclassifying these 5 GTDB species as a single species (**Fig. 3**). E. sp005843885 and E. sp004211955 also form a cluster which is unsurprising as the ANI between the GTDB representative genomes of these species is 94.95%.

**Figure 3.**
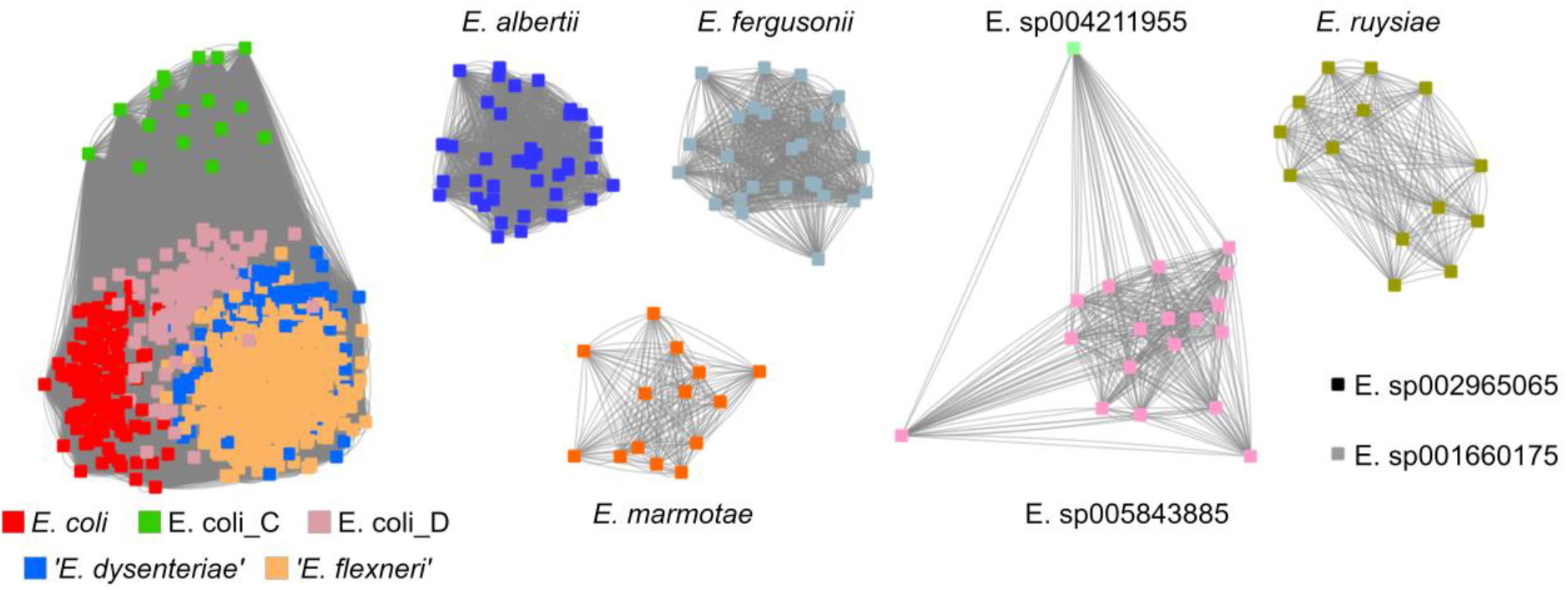
Clustering of 1,148 GTDB R06-RS202 *Escherichia* genomes dereplicated at 99% ANI. Each node represents a genome and two genomes are connected by an edge if their Mash distance is ≤0.05, which is approximately equivalent to an ANI ≥95%. Nodes are colored by GTDB species assignment. The graph layout was determined using the force directed method implemented in Cytoscape (Shannon et al., 2003).

## Conclusions

In previous releases of GTDB, we attempted to retain at least some *Shigella* species as distinct entities within the genus *Escherichia, i*.*e. ‘E. flexneri’* and *‘E. dysenteriae’*, using our type strain nucleated ANI-based species delineation approach (Parks et al., 2020), albeit with reduced ANI radii (**Fig. 1A**). This was a compromise to accommodate the well-known issue of *Shigella* species being genomically closely related to *E. coli* (Brenner et al., 1973; Richter and Rosselló-Móra 2009; Jain et al., 2018; Ciufo et al., 2018), while still retaining some of the clinically important classification information associated with *Shigella*. An unforeseen consequence of this compromise was the reclassification of almost 60% of *E. coli* genomes including *E. coli* K-12 to ‘*E. flexneri’* (**Table 1**), which is a source of potential confusion. Indeed, K-12 is often mistakenly thought to be the type strain of *E. coli*, but in fact is genomically and phenotypically quite distinct from the actual type strain, *E. coli* U5/41^T^ (DSM 30083^T^), which has been sequenced only comparatively recently (Meier-Kolthoff et al., 2014). To remove this confusion, we intend to treat *Shigella* species as later heterotypic synonyms of *E. coli* (**Fig. 1B**) in the next GTDB release, consistent with previous genomic circumscriptions (Lan and Reeves, 2002; Richter and Rosselló-Móra, 2009; van den Beld and Reubsaet, 2012), which also returns strain K-12 to the species *E. coli*. The other major consequence of this change is that the GTDB taxonomy as it currently stands is not useful for clinical classification of *Shigella* isolates. In our opinion, reclassifying *Shigella spp*. as later heterotypic synonyms of *E. coli* will ultimately best serve the community as this adheres to the species definition adopted by the GTDB, makes *E. coli* commensurate with the majority of bacterial species, and most accurately reflects the phylogenetic relationship and genomic similarity of *E. coli* and *Shigella* strains.

## Methods

### *Escherichia/Shigella* genome dataset

The 23,686 genomes classified as *Escherichia* in GTDB R06-RS202 or *Escherichia/Shigella* according to the NCBI Taxonomy (Schoch et al. 2020; downloaded September 23, 2020) were considered in this study. Two notable exceptions are inclusion of the *E. ruysiae* type strain genome GCF_902498915.1 which was released after GTDB R06-RS202 and filtering of the genome GCF_009711095.1 which is classified as ‘*E. alba’* at NCBI but recognized as belonging to the genus *Intestinirhabdus* (Xu et al., 2020). The *E. ruysiae* type strain genome (GCF_902498915.1) was compared to GTDB species representatives in *Escherichia* and found to be highly similar to the representative of Escherichia sp000208585 (GCF_000208585.1) with an ANI of 99.8% and AF of 0.97. Consequently, the 32 genomes in the GTDB R06-RS202 species cluster Escherichia sp000208585 are referred to as *E. ruysiae* in this study. A high-quality set of 23,538 genomes defined as having a completeness ≥90% and contamination ≤5% as estimated using CheckM v1.1.3 (Parks et al., 2015) was used for analyses that might be negatively impacted by the presence of lower-quality genome assemblies.

### Inferring core gene tree

Prokka v1.14.6 (Seemann, 2014) using the *Escherichia* specific database was used to annotate genomes and the core gene set determined using Roary v3.12.0 (Page et al., 2015). The core gene set for the 16 *Escherichia/Shigella* type strain or GTDB representative genomes was defined as any gene present in ≥14 of the genomes which resulted in a set of 2,329 core genes. These genes were aligned with MAFFT v7.394 (Katoh et al., 2013) which produced a 2,263,396 base multiple sequence alignment. The tree was inferred with IQ-Tree v1.6.12 (Nguyen et al., 2015) using ModelFinder (Kalyaanamoorthy et al., 2017) to establish GTR+F+R5 as the best-fit model and tree stability assessed with the SH-aLRT test and ultrafast bootstraps set to 1,000 replicates (Hoang et al., 2018). The tree was rooted on *E. albertii* based on a preliminary tree using *Salmonella enterica* LT^T^ (GCF_000006945.2) as an outgroup which indicate this *Escherchia* species to be the most basal member of the genus. This rooting is also in agreement with *E. albertii* being the most basal species in the ANI-based UPGMA tree.

### Calculating ANI and AF

The ANI and AF between genomes was calculated with FastANI v1.3 (Jain et al., 2018) with default parameters unless otherwise indicated. Since the ANI and AF values produced by FastANI are not symmetric, the maximum of the two reciprocal calculations was used in agreement with the approach adopted by GTDB (Parks et al., 2020). ANI values were also calculated with the *ani*.*rb* script in the Enveomics Collection (Rodriguez-R LM & Konstantinidis, 2016) using default parameters and BLASTn 2.9.0+ (Camacho et al., 2009) to identify orthologous fragments.

Mash v2.0 (Ondov et al., 2016) with a sketch size of 5,000 and *k*-mer size of 16 was used to estimate the ANI between all 554,013,906 pairwise combinations of the 23,538 high-quality GTDB R06-RS202 *Escherichia* genomes as the use of less computationally efficient methods was not practical. Genomes were dereplicated in a greedy manner consisting of three steps: i) sort genomes in descending order of estimated assembly quality, ii) select the highest-quality genome to form a new cluster, and iii) assign any unclustered genome with a Mash distance <0.01 (≈99% ANI) to this new cluster. These steps were repeated until all genomes were assigned to a cluster. Estimated assembly quality was defined as completeness – 5*contamination as estimated using CheckM v1.1.3 (Parks et al., 2015).

Pairwise ANI values were visualized as a UPGMA tree calculated with DendroPy v4.5.1 (Sukumaran and Holder, 2010) and as a graph generated with force directed layout method implemented in Cytoscape (Shannon et al., 2003).

## Supplementary Figures and Tables

**Supplementary Figure 1.**
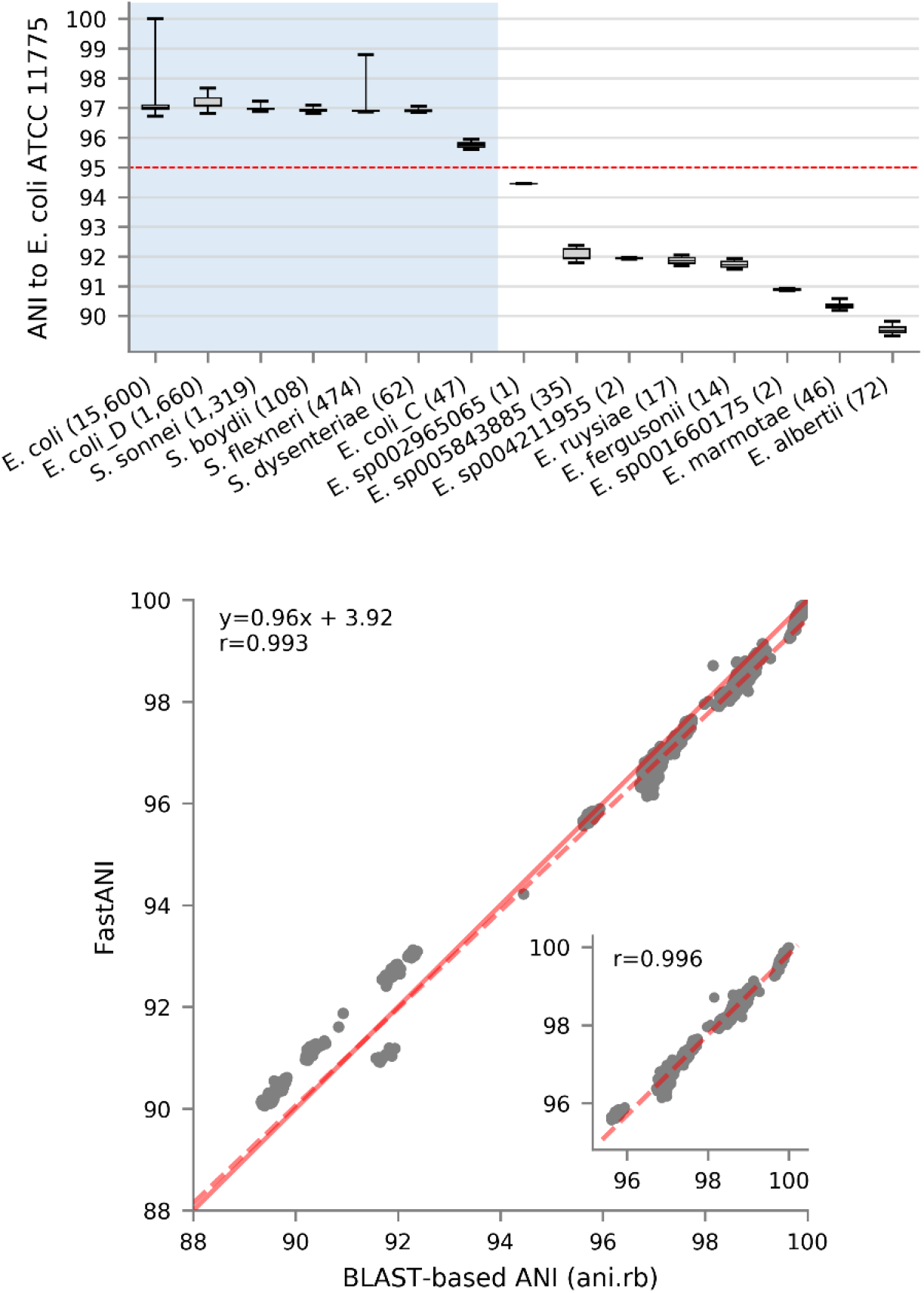
ANI between the *E. coli* ATCC 11775^T^ genome and all genomes classified as *Escherichia* in GTDB R06-RS202. (**A**) ANI values calculated with the BLAST-based *ani*.*rb* script in the Enveomics Collection. This plot is analogous to the FastANI results provided in Figure 2C. (**B**) Correlation between ANI values calculated with FastANI and *ani*.*rb*. The y=x line is shown in solid red, the best-fit line is shown as red dashes, and the Pearson correlation coefficients is given in the upper left corner. The inset plot shows results for *E. coli*, ‘*E. flexneri’*, ‘*E. dysenteriae*’, E. coli_C, and E. coli_D genomes.

**Supplementary Table 1.** Incongruent species classification between GTDB and NCBI (see Excel file).

**Supplementary Table 2.** NCBI species with the largest number of reassignments in GTDB.

| NCBI species | No. genomes | No. reassigned genomes | Reassigned genomes (%) | GTDB species |
| --- | --- | --- | --- | --- |
| *Escherichia coli* | 20,973 | 16,609 (79.19%) | 28.28 | *E. flexneri*: 12,400 (59.12%); *E. coli*: 4,364 (20.81%); *E. coli_D*: 2,094 (9.98%); *E. dysenteriae*: 2,044 (9.75%); *E. coli_C*: 45 (0.21%); *E. sp000208585*: 9 (0.04%); *E. sp005843885*: 2 (0.01%); *Klebsiella quasipneumoniae*: 1 (0.00%); *E. coli_E*: 1 (0.00%); *Phytobacter ursingii*: 1 (0.00%); *Citrobacter freundii*: 1 (0.00%); *Enterobacter roggenkampii*: 1 (0.00%); *Pseudomonas_E* sp002113295: 1 (0.00%); *E. albertii*: 1 (0.00%); *Kluyvera georgiana_A*: 1 (0.00%); *Hafnia alvei*: 1 (0.00%); *E. marmotae*: 1 (0.00%); *Citrobacter gillenii*: 1 (0.00%); *Providencia rettgeri_D*: 1 (0.00%); *Enterobacter hormaechei_A*: 1 (0.00%); *Klebsiella aerogenes*: 1 (0.00%); *Klebsiella variicola*: 1 (0.00%) |
| *Enterococcus faecium* | 2,019 | 2,019 (100.00%) | 3.44 | *Enterococcus_B* faecium: 1,793 (88.81%); *Enterococcus_B* lactis: 224 (11.09%); *Enterococcus_D* sp002850555: 2 (0.10%) |
| *Listeria monocytogenes* | 3,525 | 1,740 (49.36%) | 2.96 | *L. monocytogenes*: 1,785 (50.64%); *L. monocytogenes_B*: 1,562 (44.31%); *L. monocytogenes_C*: 172 (4.88%); *L. innocua*: 6 (0.17%) |
| *Mycobacteroides abscessus* | 1,686 | 1,686 (100.00%) | 2.87 | *Mycobacterium abscessus*: 1,686 (100.00%) |
| *Campylobacter jejuni* | 1,678 | 1,678 (100.00%) | 2.86 | *Campylobacter_D* jejuni: 1,670 (99.52%); *Campylobacter_D* lari_C: 3 (0.18%); *Campylobacter_D* jejuni_D: 2 (0.12%); *Campylobacter_D* jejuni_C: 1 (0.06%); *Campylobacter_D* jejuni_B: 1 (0.06%); *Campylobacter_D* jejuni_A: 1 (0.06%) |
| *Burkholderia pseudomallei* | 1,604 | 1,604 (100.00%) | 2.73 | *Burkholderia mallei*: 1,604 (100.00%) |
| *Pseudomonas viridiflava* | 1,518 | 1,518 (100.00%) | 2.58 | *P._E* viridiflava: 1,355 (89.26%); *P._E* avellanae: 20 (1.32%); *P._E* koreensis_C: 13 (0.86%); *P._E* orientalis_A: 12 (0.79%); *P._E* sp002979555: 9 (0.59%); *P._E* canadensis: 9 (0.59%); *P._E* sp005233515: 8 (0.53%); *P._E* congelans: 7 (0.46%); *P._E* sp000952175: 6 (0.40%); *P._E* sp002843605: 6 (0.40%); *P._E* sivasensis: 6 (0.40%); *P._E* salomonii: 5 (0.33%); *P._E* lurida: 5 (0.33%); *P._E* sp900596015: 5 (0.33%); *P._E* sp001297015: 5 (0.33%); *P._E* sp900583165: 4 (0.26%); *P._E* sp900582625: 4 (0.26%); *P._E* sp900589395: 3 (0.20%); *P._E* sp900585815: 3 (0.20%); *P._E* syringae: 3 (0.20%); *P._E* viridiflava_C: 2 (0.13%); *P._E* viridiflava_B: 2 (0.13%); *P._E* coleopterorum: 2 (0.13%); *P._E* amygdali: 2 (0.13%); *P._E* sp900573885: 2 (0.13%); *P._E* poae: 2 (0.13%); *P._E* sp900582195: 2 (0.13%); *P._E* sp002699985: 2 (0.13%); *P._E* sp900580675: 1 (0.07%); *P._E* sp900581005: 1 (0.07%); *P._E* sp900601905: 1 (0.07%); *P._E* asturiensis: 1 (0.07%); *P._E* sp900590755: 1 (0.07%); *P._E* sp900580865: 1 (0.07%); *P._E* marginalis_B: 1 (0.07%); *P._E* ovata: 1 (0.07%); *P._E* synxantha_A: 1 (0.07%); *P._E* sp900585905: 1 (0.07%); *P._E* sp900602065: 1 (0.07%); *P._E* syringae_Q: 1 (0.07%); *P._E* sp003097075: 1 (0.07%); *P._E* sp900591205: 1 (0.07%) |
| *Enterobacter hormaechei* | 1,458 | 1,444 (99.04%) | 2.46 | *E. hormaechei_A*: 1,440 (98.77%); *E. hormaechei*: 14 (0.96%); *E. chengduensis*: 1 (0.07%); *Escherichia coli*: 1 (0.07%); *E. asburiae_B*: 1 (0.07%); *E. quasihormaechei*: 1 (0.07%) |
| *Shigella sonnei* | 1,368 | 1,368 (100.00%) | 2.33 | *E. flexneri*: 1,354 (98.98%); *E. coli_D*: 13 (0.95%); *Serratia marcescens_K*: 1 (0.07%) |
| *Bacillus cereus* | 1,094 | 1,094 (100.00%) | 1.86 | *B._A* bombysepticus: 414 (37.84%); *B._A* paranthracis: 148 (13.53%); *B._A* cereus: 137 (12.52%); *B._A* nitratireducens: 69 (6.31%); *B._A* anthracis: 56 (5.12%); *B._A* thuringiensis: 35 (3.20%); *B._A* thuringiensis_S: 32 (2.93%); *B._A* tropicus: 31 (2.83%); *B._A* wiedmannii: 26 (2.38%); *B._A* mobilis: 21 (1.92%); *B._A* thuringiensis_N: 19 (1.74%); *B._A* toyonensis: 16 (1.46%); *B._A* mycoides: 15 (1.37%); *B._A* cereus_U: 9 (0.82%); *B._A* thuringiensis_K: 8 (0.73%); *B._A* cereus_S: 7 (0.64%); *B._A* cereus_K: 7 (0.64%); *B._A* paramycoides: 5 (0.46%); *B._A* albus: 5 (0.46%); *B._A* cereus_AT: 5 (0.46%); *B._A* cereus_AU: 5 (0.46%); *B._A* luti: 4 (0.37%); *B._A* sp002584985: 3 (0.27%); *B._A* pseudomycoides: 3 (0.27%); *B._A* cereus_AG: 2 (0.18%); *B._A* cereus_AK: 2 (0.18%); *B._A* cereus_AQ: 2 (0.18%); *B._A* cereus_AW: 1 (0.09%); *B._A* proteolyticus: 1 (0.09%); *B._A* cereus_AZ: 1 (0.09%); *B._A* mycoides_C: 1 (0.09%); *B._A* sp008923725: 1 (0.09%); *B._A* cereus_AV: 1 (0.09%); *B._A* mycoides_B: 1 (0.09%); *B._A* cereus_O: 1 (0.09%) |

**Supplementary Table 3.** Genomes classified as an *Escherichia* or *Shigella* species at NCBI which are misclassified based on their ANI to the type strain of the species.

| Genome ID | NCBI species | ANI to type strain genome | AF to type strain genome | GTDB species |
| --- | --- | --- | --- | --- |
| GCF_000601195.1 | *Escherichia coli* | 92.85 | 0.87 | *Escherichia* sp000208585 |
| GCF_903932005.1 | *Escherichia coli* | 92.85 | 0.83 | *Escherichia* sp005843885 |
| GCF_003145355.1 | *Escherichia coli* | 92.84 | 0.87 | *Escherichia* sp000208585 |
| GCA_013425105.1 | *Escherichia coli* | 92.83 | 0.86 | *Escherichia* sp000208585 |
| GCF_005400045.1 | *Escherichia coli* | 92.77 | 0.85 | *Escherichia* sp000208585 |
| GCF_002110245.1 | *Escherichia coli* | 92.7 | 0.86 | *Escherichia* sp000208585 |
| GCF_004745245.1 | *Escherichia coli* | 92.65 | 0.81 | *Escherichia* sp005843885 |
| GCF_000459855.1 | *Escherichia coli* | 92.58 | 0.84 | *Escherichia* sp000208585 |
| GCF_903932105.1 | *Escherichia coli* | 92.52 | 0.88 | *Escherichia* sp000208585 |
| GCF_000398885.1 | *Escherichia coli* | 92.47 | 0.85 | *Escherichia* sp000208585 |
| GCA_001630835.1 | *Escherichia coli* | 92.43 | 0.85 | *Escherichia* sp000208585 |
| GCF_011881725.1 | *Escherichia coli* | 92.22 | 0.89 | *Escherichia coli*_E |
| GCF_903932125.1 | *Escherichia coli* | 91.09 | 0.81 | *Escherichia marmotae* |
| GCF_001286085.1 | *Escherichia coli* | 90.28 | 0.8 | *Escherichia albertii* |
| GCF_900448175.1 | *Escherichia coli* | 82.35 | 0.57 | *Citrobacter gillenii* |
| GCF_003007795.1 | *Escherichia coli* | 82.23 | 0.57 | *Citrobacter freundii* |
| GCA_011008595.1 | *Escherichia coli* | 81.4 | 0.46 | *Kluyvera georgiana*_A |
| GCA_009766545.1 | *Escherichia coli* | 81.39 | 0.51 | *Enterobacter hormaechei*_A |
| GCF_006381975.1 | *Escherichia coli* | 81.33 | 0.5 | *Enterobacter roggenkampii* |
| GCA_003175335.1 | *Escherichia coli* | 81.19 | 0.44 | *Klebsiella variicola* |
| GCA_003301495.1 | *Escherichia coli* | 81.05 | 0.44 | *Klebsiella aerogenes* |
| GCA_003363055.1 | *Escherichia coli* | 80.98 | 0.44 | *Phytobacter ursingii* |
| GCA_003176195.1 | *Escherichia coli* | 80.9 | 0.42 | *Klebsiella quasipneumoniae* |
| GCF_009647995.1 | *Escherichia coli* | 0* | 0* | *Pseudomonas*_E sp002113295 |
| GCF_011008745.1 | *Escherichia coli* | 0 * | 0* | *Hafnia alvei* |
| GCF_009648115.1 | *Escherichia coli* | 0 * | 0* | *Providencia rettgeri*_D |
| GCF_902167795.1 | *Escherichia fergusonii* | 91.03 | 0.69 | '*Escherichia flexneri*' |
| GCF_001060475.1 | *Shigella boydii* | 78.85 | 0.25 | *Serratia marcescens*_I |
| GCF_001062045.1 | *Shigella boydii* | 78.82 | 0.23 | *Serratia marcescens*_I |
| GCF_001060695.1 | *Shigella boydii* | 78.73 | 0.23 | *Serratia marcescens*_I |
| GCA_001066285.1 | *Shigella sonnei* | 78.85 | 0.24 | *Serratia marcescens*_K |

\* sufficiently divergent that FastANI fails to provide an ANI or AF value

**Supplementary Table 4.** Type strain and GTDB representative genomes.

| Genome ID | GTDB species | NCBI species | Strain ID | GTDB representative | Type strain of species |
| --- | --- | --- | --- | --- | --- |
| GCF_001660175.1 | *Escherichia* sp001660175 | unclassified | B1147 | TRUE | FALSE |
| GCF_004211955.1 | *Escherichia* sp004211955 | unclassified | E1V33 | TRUE | FALSE |
| GCF_002965065.1 | *Escherichia* sp002965065 | unclassified | MOD1-EC7003 | TRUE | FALSE |
| GCF_005843885.1 | *Escherichia* sp005843885 | unclassified | E4742 | TRUE | FALSE |
| GCF_000759775.1 | *Escherichia albertii* | *Escherichia albertii* | NBRC 107761 | TRUE | TRUE |
| GCF_011881725.1 | *Escherichia coli*_E | *Escherichia coli* | SCPM-O-B-8794 | TRUE | FALSE |
| GCF_000026325.1 | *Escherichia coli*_D | *Escherichia coli* | UMN026 | TRUE | FALSE |
| GCA_003018335.1 | *Escherichia coli*_C | *Escherichia coli* | 2013C-4282 | TRUE | FALSE |
| GCF_003697165.2 | *Escherichia coli* | *Escherichia coli* | ATCC 11775 | TRUE | TRUE |
| GCF_000026225.1 | *Escherichia fergusonii* | *Escherichia fergusonii* | ATCC 35469 | TRUE | TRUE |
| GCF_002900365.1 | *Escherichia marmotae* | *Escherichia marmotae* | HT073016 | TRUE | TRUE |
| GCF_902498915.1 | *Escherichia ruysiae* | *Escherichia ruysiae* | OPT1704 | n/a* | TRUE |
| GCF_002946735.1 | '*Escherichia flexneri*' | *Shigella boydii* | ATCC 8700 | FALSE | TRUE |
| GCF_002949675.1 | '*Escherichia dysenteriae*' | *Shigella dysenteriae* | ATCC 13313 | TRUE | TRUE |
| GCF_002950215.1 | '*Escherichia flexneri*' | *Shigella flexneri* | ATCC 29903 | TRUE | TRUE |
| GCA_002950395.1 | '*Escherichia flexneri*' | *Shigella sonnei* | ATCC 29930 | FALSE | TRUE |

\* released after GTDB R06-RS202

**Supplementary Table 5.** ANI (lower triangle) and AF (upper triangle) between type strain genomes from *E. coli* and *Shigella* species and GTDB representative genomes for E. coli_C and E. coli_D.

|  | *E. coli* | *E. coli_C* | *E. coli_D* | *S. boydii* | *S. dysenteriae* | *S. flexneri* | *S. sonnei* |
| --- | --- | --- | --- | --- | --- | --- | --- |
| *E. coli* | - | 0.84 | 0.86 | 0.80 | 0.78 | 0.78 | 0.79 |
| *E. coli_C* | 95.64 | - | 0.88 | 0.83 | 0.81 | 0.80 | 0.81 |
| *E. coli_D* | 96.81 | 96.13 | - | 0.83 | 0.81 | 0.80 | 0.81 |
| *S. boydii* | 96.47 | 95.45 | 96.95 | - | 0.78 | 0.84 | 0.86 |
| *S. dysenteriae* | 96.30 | 95.27 | 96.65 | 97.06 | - | 0.81 | 0.80 |
| *S. flexneri* | 96.40 | 95.61 | 96.95 | 97.81 | 97.00 | - | 0.80 |
| *S. sonnei* | 96.55 | 95.59 | 97.02 | 98.14 | 97.15 | 97.94 | - |

